# Flor yeasts rewire the central carbon metabolism during wine alcoholic fermentation

**DOI:** 10.1101/2021.02.27.433177

**Authors:** Emilien Peltier, Charlotte Vion, Omar Abou Saada, Anne Friedrich, Joseph Schacherer, Philippe Marullo

## Abstract

The identification of natural allelic variations controlling quantitative traits could contribute to decipher metabolic adaptation mechanisms within different populations of the same species. Such variations could result from man-mediated selection pressures and participate to the domestication. In this study, the genetic causes of the phenotypic variability of the central carbon metabolism *Saccharomyces cerevisiae* were investigated in the context of the enological fermentation. Carbon dioxide and glycerol production as well as malic acid consumption modulate the fermentation yield revealing a high level of genetic complexity. Their genetic determinism was found out by a multi environment QTL mapping approach allowing the identification of 14 quantitative trait loci from which 8 of them were validated down to the gene level by genetic engineering. Most of the validated genes had allelic variations involving flor yeast specific alleles. Those alleles were brought in the offspring by one parental strain that is closely related to the flor yeast genetic group while the second parental strain is part of the wine group. The causative genes identified are functionally linked to quantitative proteomic variations that would explain divergent metabolic features of wine and flor yeasts involving the tricarboxylic acid cycle (TCA), the glyoxylate shunt and the homeostasis of proton and redox cofactors. Overall, this work led to the identification of genetic factors that are hallmarks of adaptive divergence between flor yeast and wine yeast in the wine biotope. These alleles can also be used in the context of yeast selection to improve oenological traits linked to fermentation yield.

## Introduction

Deciphering how the considerable phenotypic diversity observed at the species level is controlled by genetic variation is an important and non-trivial goal in biology. Improving knowledge regarding genotype-phenotype relationship provides information on evolution and adaptation mechanisms (Olson-Manning, Wagner, and Mitchell-Olds 2012) and is precious in many biological fields like medicine (Minikel et al. 2020) or food industry (McCouch 2004; Marullo et al. 2006; Sharmaa et al. 2015). Unravelling the genetic basis of adaptation highlights how organisms adapt to new selection pressure like climate change, new pathogens or drugs and vaccines (Olson-Manning, Wagner, and Mitchell-Olds 2012; Alföldi and Lindblad-Toh 2013). Domestication is a specific case of adaptation with important phenotypic change emerging from human artificial selection. Domesticated organisms are a great opportunity to study adaptation as there is a better knowledge of their adaptive history through their well-characterized phenotypic properties and selective environments (Ross-Ibarra, Morrell, and Gaut 2007; Gladieux et al. 2014). The identification of genes and molecular mechanisms leading to adaptation against domestication is also very useful in genetic selection in order to improve traits of economic interest and bringing phenotypic novelty to domesticated species (McCouch 2004).

The yeast *Saccharomyces cerevisiae* rapidly emerged as an excellent model to study genotype-phenotype relationship (Steinmetz et al. 2002; Brem et al. 2002) and plenty of quantitative genetic studies were carried out in this species to study epistasis (Sinha et al. 2006), missing heritability (Bloom et al. 2013), gene-environment interaction (Smith and Kruglyak 2008; Bhatia et al. 2014; Yadav, Dhole, and Sinha 2016; Peltier et al. 2018) or impact of rare variants (Fournier et al. 2019; Bloom et al. 2019). *S. cerevisiae* was subjected to multiple domestication events in association with a large number of human associated environments (wine, beer, bread etc.) leading to distinct phylogenetic groups (Peter et al. 2018; Sicard and Legras 2011; J. L. Legras et al. 2018). Several genetic marks of adaptation were identified such as gene loss of function (Will et al. 2010), translocations (Zimmer et al. 2014; Pérez-Ortín et al. 2002), introgressions (Novo et al. 2009; Marsit et al. 2015), and SNPs (Peltier et al. 2019) (see for review: (Giannakou, Cotterrell, and Delneri 2020). Flor and wine yeasts are both associated with wine making environment and form two distinct but closely related phylogenetic groups (J. L. Legras et al. 2018). While both groups are able to efficiently perform wine fermentation, flor yeasts used in Sherry-like wines have the specific ability to shift to oxidative metabolism and form a velum covering wine surface after fermentation (J. Legras et al. 2016). Differences in genomic content between wine and flor yeast were observed and the impact of allelic variations involved in biofilm formation were proposed as a feature of genetic adaptation (Fidalgo et al. 2006; Coi et al. 2017). Other functional adaptation hallmarks related to active gluconeogenesis, response to osmotic pressure and metal transport were predicted by a population genomic approach but have not been demonstrated yet at the gene level (Coi et al. 2017).

Recent global warming caused the steady increase of sugar content in grape juices leading to higher ethanol concentration in wine with several issues regarding consumer health and wine quality (Dariusz R. Kutyna et al. 2010). Therefore, there is a growing demand for the development of new technologies to reduce alcohol content in wine. In this context, several institutions have attempted a biological approach in order to select new strains of *S. cerevisiae* with a lower fermentation yield. Various strategies were implemented such as adaptive evolution (Tilloy et al. 2015; D. R. Kutyna et al. 2012), interspecific breeding (da Silva et al. 2015), and genetic engineering (Rossouw et al. 2013; Ehsani et al. 2009). Here, we aim at finding out undescribed natural genetic variations controlling the central carbon metabolism in order to modulate the efficiency of sugar into ethanol conversion (Fermentation yield). By applying a Quantitative Trait Loci (QTL) mapping approach, we investigated the genetic determinism of three traits (glycerol production, CO_2_ production and malic acid consumption) that shape the carbon balance in enological conditions.

Our study is based on the analysis of a progeny obtained by crossing two strains derived from wine starters. A deeper analysis of parental genomes showed that, unexpectedly, one of the parental strains results to have a mosaic genome inherited from both wine and flor yeasts while the second parental strain belongs to the wine group. This admixture has promoted an important phenotypic variability impacting the central carbon metabolism of the F1 progeny. A total of 14 QTLs were identified and the effect of eight of them were experimentally validated down to the gene level. Six genes (*PMA1*, *PNC1*, *PYC2*, *SDH2, MAE1,* and *MSB2*), among which three are directly involved in central carbon metabolism (*SDH2* in tricarboxylic acid cycle (TCA)), *MAE1* in pyruvate metabolism and *PYC2* in gluconeogenesis pathways, show allelic variations highly specific to flor yeasts group. Linked to these validated genes, further proteomic analyses highlighted different metabolic regulations between the parental strains for TCA and glyoxylate shunt. Altogether, these results support the hypothesis that allelic variations between wine and flor yeasts generate important phenotypic differences and could be considered as hallmarks of adaptation for different growth strategies on the wine biotope. These results also show that flor yeasts constitute a great reservoir of genetic variation to bring phenotypic novelty in commercial yeast starter to cope for new challenges as global warming (Mira de Orduña 2010) and new viticultural practices (Kontoudakis et al. 2011).

## Results

### Biometric study of the glycerol, CO*2* and malic acid

In order to explore the genetic determinism of central carbon metabolism during wine alcoholic fermentation, the previous dataset of fermentation traits measured within a QTL mapping population was used (Peltier, Sharma, et al. 2018). This population was obtained by mating two fully homozygous strains (SB and GN) derived from the sporulation of wine starters. A total of 94 meiotic segregants were obtained though sporulation of a single hybrid (SBxGN) (Fig1) and phenotyped in three environmental conditions using a small-scale fermentation dispositive and enzymatic assays to measure fermentation kinetics traits and endpoint concentration of several metabolites, including glycerol and CO_2_ production. All segregants were sequenced and a genetic map of 3433 biallelic markers was built in order to identify the genetic factors controlling these phenotypes (Table S1). In the present study, an additional phenotyping effort was achieved by measuring malic acid consumption in the same conditions.

**Figure 1.**
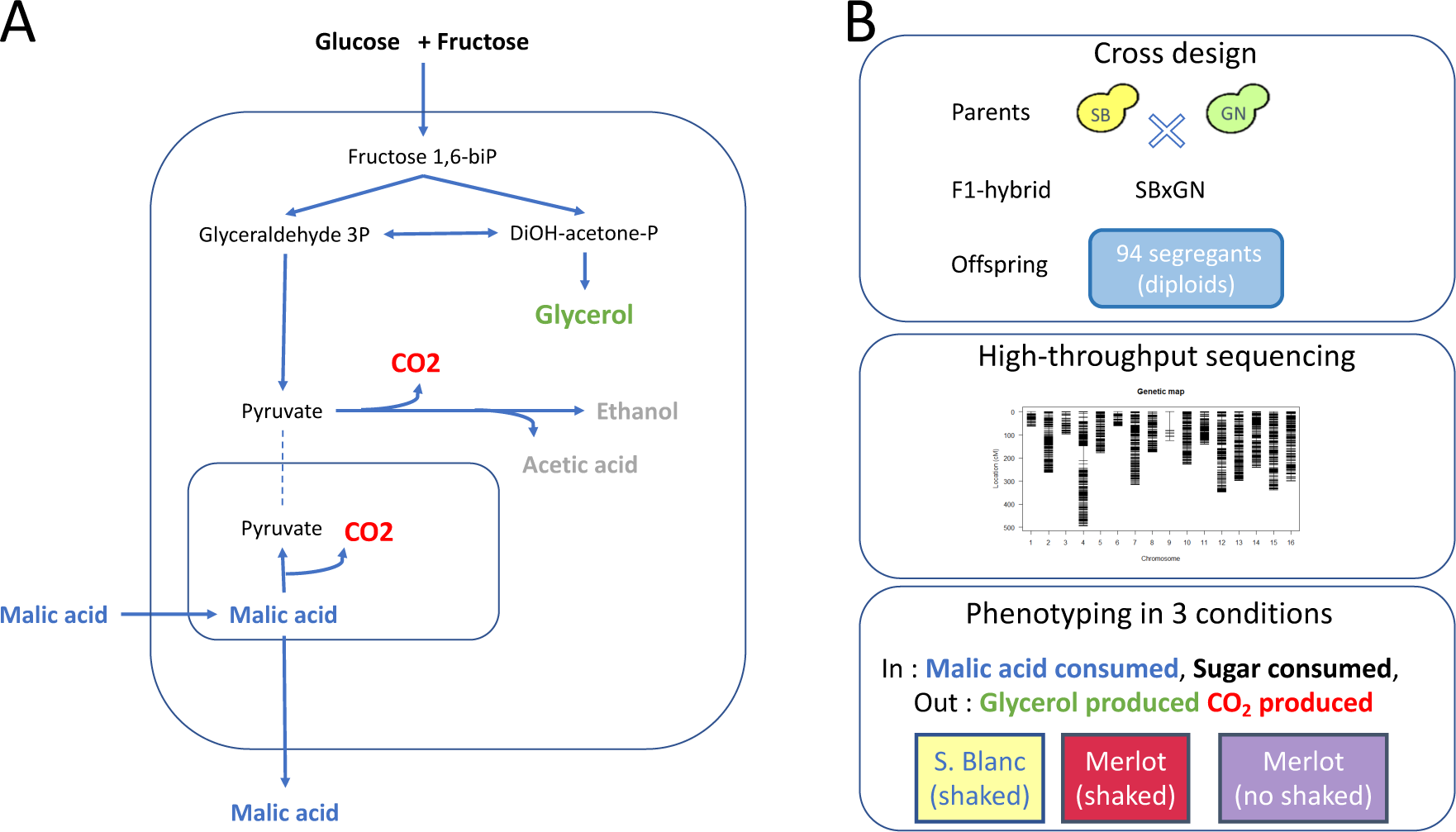
Experimental design. **Panel A.** Overview of yeast central carbon metabolism during fermentation with the main carbon input and output. **Panel B.** Segregant population, genetic map and phenotypic conditions used for QTL mapping.

Carbon balance was evaluated by measuring the main organic compounds assimilated and/or produced for each of the 94 segregants at the end of the alcoholic fermentation (Table S2). According to the must, the fermentation yield computed ranged between 0.45 and 0.48 which is close to values observed in other studies (Tilloy, Ortiz-Julien, and Dequin 2014) (Supplementary file S1). An analysis of variance demonstrated a significant genetic (strain) impact on the fermentation yield (17% of the total variance explained). This integrative trait is mostly shaped by the quantitative variation of three metabolites: glycerol, malic acid, and CO_2_ that were partially correlated (Figure S1). Glycerol and CO_2_ (which is stoichiometrically linked to ethanol) are *de novo* synthetized by yeast catabolism; their concentrations are expressed in g/L. The final concentration of CO_2_ produced is expressed hereafter as *CO_2_max*. The final concentration of malic acid depends on its initial amount in grape must which differs according to the grape juice. Since this organic acid is partially metabolized by yeast, the strain contribution was normalized by computing the percentage of Malic Acid Consumed (*MAC%*). For each trait, parental strains SB and GN are significantly different with important gaps for glycerol and *MAC%* (Wilcoxon test, pval <0.05). Indeed, SB produces 1.6 g/L more glycerol (+30%) and consumes 28% more malic acid than GN. Since malic consumption and glycerol production have an opposite effect on CO_2_ and ethanol production, the phenotypic differences for CO_2_ are sharper. These differences are consistent with previous results showing that SB is the top strain for glycerol production and malic acid consumption compared to a panel of commercial starters (Peltier, Bernard, et al. 2018).

Each trait had a high overall heritability (Table S3) and displayed a bell-shaped distribution with number of segregants showing transgressive values respect to parental strains (Fig S2). These broad biometric observations highlighted a polygenetic control of each trait with a positive contribution of both parental strains.

### Linkage analysis brings out a linkage hotspot with pleiotropic effect

In a previous work that explored QTL interaction with environment, five QTLs were associated with *CO_2_max* and *glycerol* production in the SBxGN offspring (Peltier *et al*., 2018). Here, we aimed at identifying supplemental QTL controlling *MAC%* that was newly phenotyped. A linkage analysis was performed and significantly associated nine QTLs to this trait. Therefore, a total of 14 QTL are involved in *CO_2_max, glycerol* and *MAC%* (Fig 2 and Table S4). The effects of parental alleles are shown in the Fig S3. Intriguingly, a large region of the chromosome VII (387 kb to 716 kb) was associated with all the considered traits. This linkage hotspot is almost entirely above the significance threshold for at least one trait and four distinct linkage peaks can be distinguished. This hotspot encompasses one major QTL, the locus *VII_415* (Chr VII, position 415,719), influencing the glycerol production (LOD score >10) which explains more than 10 % of total variance. Interestingly, for this cross, a sharper region of chromosome VII (50 kb) was previously associated with kinetic traits during second fermentation of sparkling wines (Martí-Raga et al. 2017). Three genes of this large QTL (*PDR1*, *PMA1* and *MSB2*) were demonstrated to have an important phenotypic impact in this condition. Here, the QTL *VII_482* linked to *MAC%* is located in the *PMA1* coding sequence (479,910 482,666).

**Figure 2.**
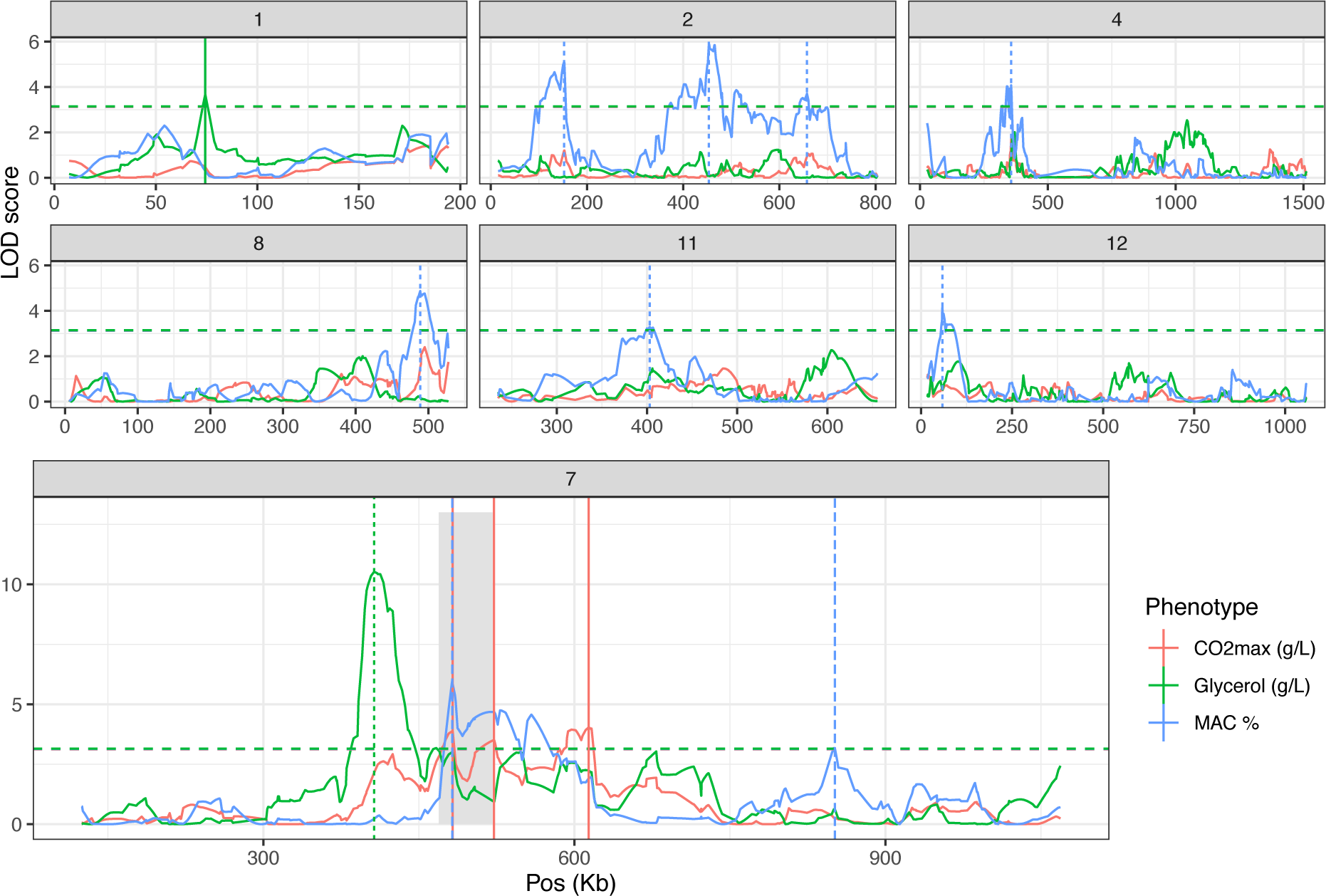
Linkage analysis leads to the identification of 14 QTLs. Linkage analysis results for the *CO_2_max*, *Glycerol* and *MAC%* for chromosome with at least one QTL. Horizontal lines represent the threshold of significance according to permutation test (FDR = 5 %). Vertical lines highlight QTL peaks. Grey shadow encompasses the previously identified QTL hotspot containing *PDR1*, *MSB2* and *PMA1* (Martí-Raga *et al*., 2017).

### Multiple Quantitative Trait Genes control glycerol production and malic acid consumption

Candidate genes neighboring the QTL peak within a 20 kb window were considered through their functional annotation and by checking for ns-SNPs within parental strains sequences using the algorithm SnpEff (Table S5) (Sherman and Salzberg 2020). We selected also the three genes (*PDR1*, *PMA1* and *MSB2*) previously validated for second fermentation traits that are located near the major hotspot of chromosome VII in the present work. This leads to consider 11 candidate genes that could impact the traits investigated. Their effects were interrogated by a Reciprocal Hemizygosity Analysis (RHA) (Steinmetz et al. 2002). The impact of parental alleles was compared in alcoholic fermentation test using the same fermentation protocol. In addition, ethanol content (% Vol) was estimated by infrared reflectance rather than enzymatic assay (see methods). The effect of four candidate genes impacting *CO_2_max* and/or *glycerol* was tested. They belong to the two major QTLs found in term of variance explained: *ADE6* (*VII_616*), *MSB2* (*VII_512*), *PDR1* (*VII_482*), *PNC1* (*VII_415*). The RHA was carried out in the M15_sk condition with two sugar concentration levels (219 and 265 g/L) using at least five biological replicates for each condition. Sugar spiking would emphasize the phenotypic differences related to CO_2_ and ethanol production. The most obvious effects were obtained for glycerol production for genes *ADE6*, *MSB2*, and *PCN1* for which hemizygous hybrids are significantly different (Wilcoxon test, pval < 0.1) (Fig 3, panel A). These three genes are located in a region of 200 kb along the chromosome VII hotspot demonstrating that distinct genetic factors in this region control the glycerol production.

**Figure 3.**
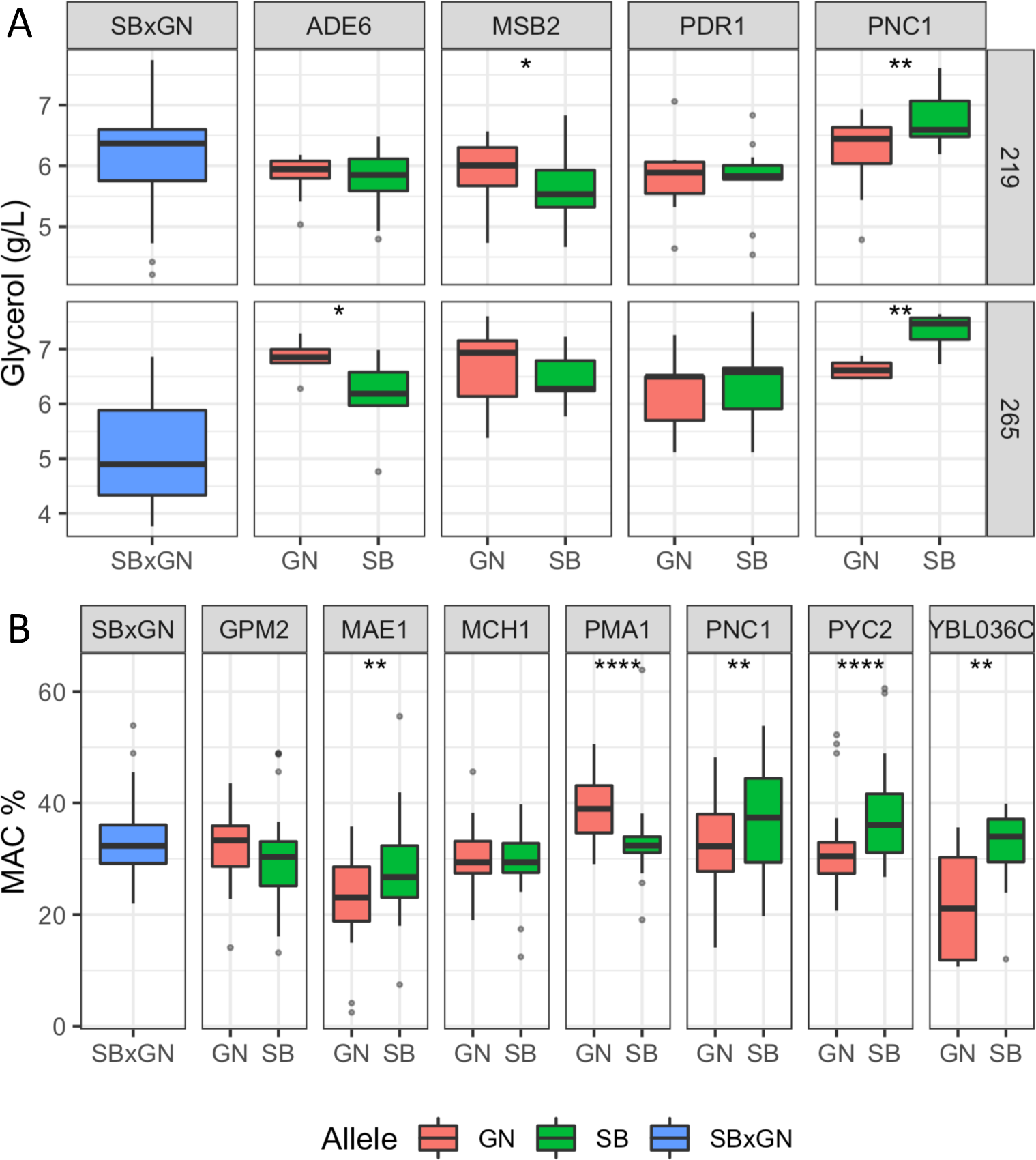
Results of the reciprocal hemizygosity analysis. Boxplot are colored according to the allele present in the hemizygous hybrids (blue = both, red = GN and green = SB) and represented the dispersion of at least five biological replicates. A Wilcoxon–Mann–Whitney test was applied to assess the significance of the phenotypic difference between hemizygotes. The level of significance is indicated as follows: * *p* ≤ 0.1, ** *p* ≤ 0.05, *** *p* ≤ 0.01 and *p* ≤ 0.001****. Panel A. RHA result for glycerol. Panel B. RHA result for *MAC%*.

Intriguingly, the sugar content modulated the phenotypic responses of hemizygous hybrids. Indeed, in sugar-spiked grape must (M15_265), alleles *ADE6*^GN^ enhanced glycerol production of 12 %, while the allele *MSB2^GN^* has an enhancer effect only in the original M15 grape must (219g/L of initial sugar). The allelic forms *ADE6^GN^*, *PNC1^SB^* promote the glycerol production and their effects are those observed in the SBxGN progeny (Table S4, Fig S3). In contrast, the *MSB2^GN^* allele produced more glycerol which is not observed in the segregating progeny (Fig S4). This opposite effect has been previously described for the same gene for another phenotype and could be due to the complex genetic architecture of chromosome VII (Martí-Raga et al. 2017). The difference observed in glycerol production for *ADE6*, *PNC1* and *MSB2* did not impact either the *CO_2_max* or the ethanol content.

In the same way, seven candidate genes belonging to six QTLs affecting *MAC%* were evaluated: *MAE1* (*XI_381*), *MCH1* and *GPM2* (*IV_356*), *PYC2* (*II_669*), *PMA1* (*VII_482*), *SDH2* (*XII_53*) and *YBL036c* (*II_152*). Fermentations were carried out in both M15 and SB14. RHA revealed a significant effect for the genes *MAE1*, *PMA1, PYC2* and *YBL036c* (Fig 3, panel B) (Wilcoxon test, pval < 0.05). The alleles of *MAE1*, *PYC2* and *YBL036c* inherited from the parental strain SB consumed respectively 25%, 19%, and 45% more malic acid than those inherited from GN. In contrast, the *PMA1^GN^* allele consumed 18% more malic acid than *PMA1^SB^*. This gene, encoding for the plasma membrane ATPase, has been previously linked to the maintenance of pH homeostasis during wine fermentation and is located in the center of chromosome VII hotspot (Martí-Raga et al. 2017). Unexpectedly, a significant effect of *PNC1* on *MAC%* was also observed and the hemizygote hybrid harboring the *PNC1^SB^* allele consumes 15 % more malic acid than *PNC1*^GN^ (Fig 3, panel B) (Wilcoxon test, pval < 0.05). The genomic position of *PNC1* is about 50 kb from the nearest QTL peak for *MAC%* VII_482), however the other causative genes (*PMA1*, *MSB2, ADE6*) associated with the chromosome VII hotspot may have altered the precision of our linkage analysis.

Beside the validation of these five genes on *MAC%*, reciprocal hemizygous analysis of *SDH2* suggested its potential contribution on malic acid consumption. Although the hemizygous are not statistically different, a strong haploinsufficiency effect in both hemizygous hybrids was observed affecting either *MAC%* (−14%) and fermentation kinetics by doubling the fermentation duration (Fig S5). Intriguingly, this haploinsufficiency was only present in M15 grape juice. Two factors suspected to have an impact on this haploinsufficiency were tested (initial malic concentration and pH) in synthetic grape juice (SGJ) by adjusting these two initial values to either M15 or SB14 levels. An haploinsufficiency similar to that in M15 was found in all four conditions even in the one mimicking SB14 conditions (Fig S5). No significant interaction between the level of haploinsufficiency and pH and malic acid was found (Anova, pval > 0.1). These findings suggest that *SDH2* has a great impact on fermentation rate and *MAC%* during grape juice fermentation. However, since the RHA test was limited by the haploinsufficiency effect our experiments failed to clearly demonstrate the impact of parental allelic variations.

Altogether, these functional analyses validated the role of eight Quantitative Trait Gene (QTG). Four of them play a direct role in the central metabolism encoding enzymes involved in oxidoreductive reactions of carbohydrate metabolism (*MAE1*, *PYC2*, *PNC1*, *SDH2*). Two others are key regulators of osmotic (*MSB2*) and pH (*PMA1*) homeostasis. The RHA also revealed that *ADE6* and *YBL036c* contribute to the phenotypic difference between the parental strains for glycerol production and malic acid consumption, respectively (Fig 3). However, their functional connection with the metabolic pathway of glycerol and malic acid is more difficult to address at this stage.

### SB is a mosaic strain derived from flor and wine yeasts

QTL mapping is a useful strategy for identifying natural genetic variations that shape phenotypic diversity between two strains. However, in most of the cases, the causative mutations identified are rare and specific to one parental strain (Bloom et al. 2019; Fournier et al. 2019; Peltier et al. 2019) due to the clonal structure of *S. cerevisiae* population (Peter et al. 2018). This impairs the identification of more general mechanisms of adaptation resulting to natural selection. In order to have a more precise idea of the evolutive relevance of QTL identified, SB and GN genomes were compared to those of 403 wine related strains previously released (Peter et al. 2018; Legras et al. 2018). A phylogenetic tree was generated using 385,678 SNPs discriminating the 403 wine strains plus the parental strains SB and GN. This collection of strains encompasses wine (n=358) and flor (n=47) strains that form distinct groups as previously described (Coi et al. 2017; Legras et al. 2018) (Table S6). Interestingly, SB is genetically close to the flor group while GN is quite similar to the wine group (Fig 4, panel A). Consequently, the two parental strains used in this study are quite distant with a sequence divergence of 0.19 % (∼22,000 SNPs). The relatedness of SB genome with the flor group was deeply investigated by selecting a subset of 5,086 SNPs highly specific to the flor yeast group. Those SNPs have a frequency difference higher than 90 % between flor and wine yeast groups. The strain SB harbors 44.3 % of flor yeast specific alleles while GN only has 1.7 % of them. Their distribution across the SB genome is not uniform (Fig 4, panel B). Indeed, long portions of chromosomes have inherited 100 % flor-specific alleles (Chr II) while other portions are totally exempt of them (Chr VIII). This analysis demonstrated that SB is a mosaic strain between wine yeast and flor yeast, a feature shared with some others wine starters (Coi *et al*., 2017).

**Figure 4.**
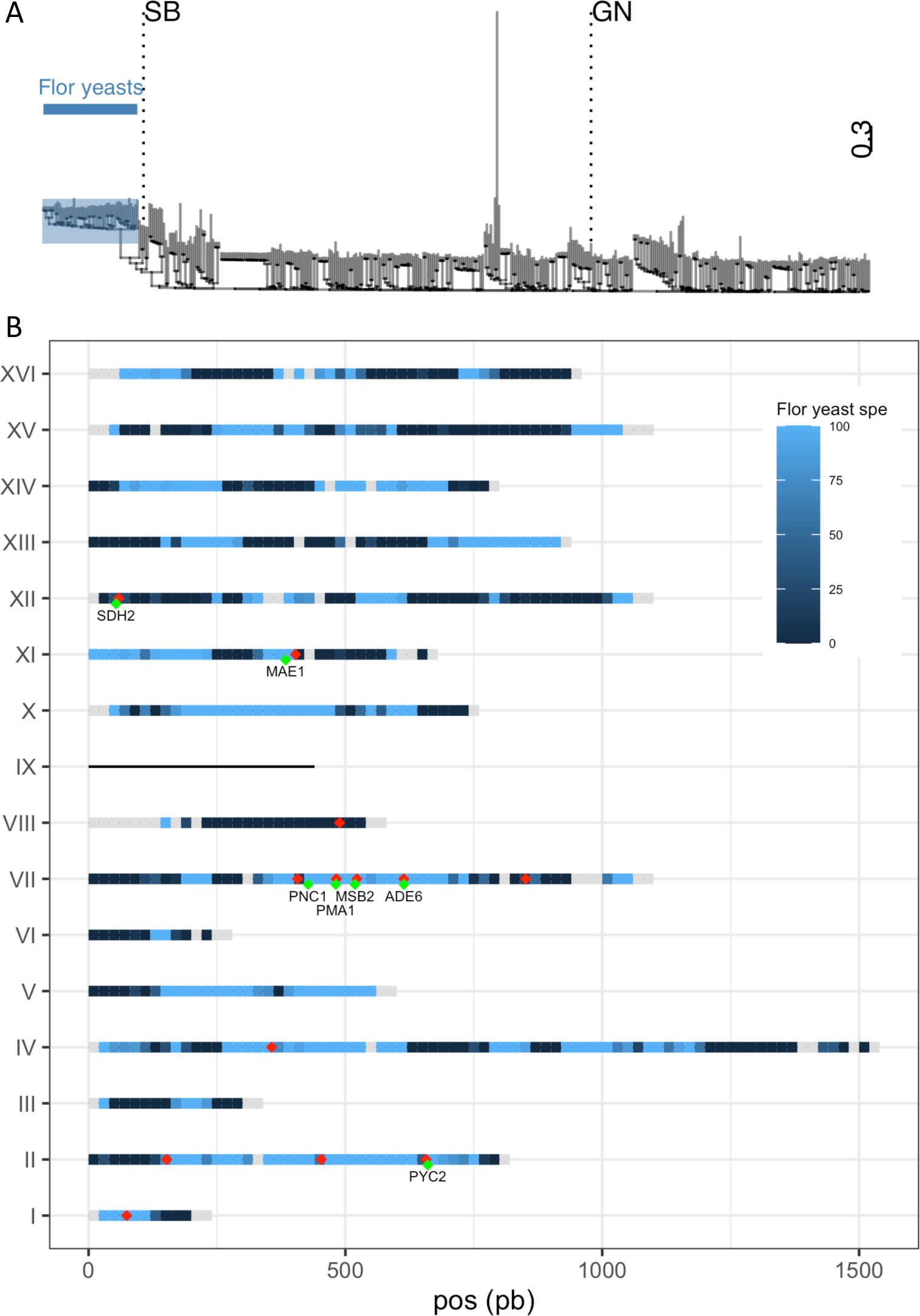
SB is closely related to flor yeasts. **Panel A.** Dendrogram using 385,678 SNPs from 405 wine strains. Flor yeasts group is highlighted. **Panel B.** Percentage of specific allele own by SB along the genome is represented by a gradient from dark blue (0 %) to light blue (100 %). Grey portions represent genome tracks without any flor yeast specific allele. SB is aneuploid for chromosome IX and therefore is not considered in this analysis. The 20 QTLs mapped are shown with red dots (some of them are overlapping) and validated genes are shown in green.

Intriguingly, nine of the fourteen QTLs mapped are located in flor specific chromosomic portions. This is the case of a large stretch within chromosome VII encompassing four causative genes (*PNC1*, *MSB2*, *PMA1*, *ADE6*) that displays the genomic signature of flor yeasts. A similar observation can be made for chromosome II in which three QTLs were identified (Fig 4, panel B). During their domestication, flor yeasts accumulated numerous mutations leading to an adaptation to grow on wine surface (Coi et al. 2017). In order to narrow such natural genetic variations, we listed the pool of ns-SNP discriminating SB and GN in the sequence of causative genes. For those SNPs, allelic frequencies of flor and wine groups were computed (Table 1). In *ADE6*, ns-SNPs listed are scarcely found whatever the group. The low allelic frequency of such polymorphisms would reflect recent mutations which is a common feature of the *S. cerevisiae* population. In contrast, for the other genes *PMA1*, *PNC1*, *PYC2*, *SDH2, MAE1,* and *MSB2*, the SB alleles are highly specific to flor yeast group while GN alleles are specific to the wine group. Therefore, these flor-specific alleles would have promoted the wide phenotypic variability of carbon metabolism observed in SBxGN progeny and more broadly are explaining phenotypic differences between flor and wine yeasts.

**Table 1.** ns-SNPs in validated genes according to genetic group.

| ORF | Gene | Protein size | Trait impacted | ns-SNP |  | Frequency in |  | deleterious effect<sup>a</sup> |
| --- | --- | --- | --- | --- | --- | --- | --- | --- |
|  |  |  |  | Protein allele | Inheritance | wine group | flor group |  |
| YGR061C | ADE6 | 1359 | Glycerol | F181L | SB | 1.7 | 12.8 | no |
|  |  |  |  | V570I | SB | 1.8 | 3.2 | no |
|  |  |  |  | P745S | GN | 0.4 | 0 | no |
|  |  |  |  | V1238A | SB | 2 | 4.3 | yes |
| YGL008C | PMA1 | 919 | MAC% | P74L | GN | 96.3 | 0 | yes |
|  |  |  |  | L176M | SB | 0.6 | 27.7 | no |
|  |  |  |  | D200E | SB | 0.6 | 10.6 | yes |
|  |  |  |  | E283R | SB | 2.7 | 100 | no |
|  |  |  |  | L290V | SB | 0.6 | 27.7 | no |
|  |  |  |  | K431I | SB | 0 | 0 | no |
|  |  |  |  | Q432E | SB | 0 | 0 | no |
|  |  |  |  | D718N | SB | 3.5 | 100 | no |
|  |  |  |  | E875Q | SB | 2.7 | 97.9 | yes |
| YGL037C | PNC1 | 217 | Glycerol. MAC% | V112A | SB | 1.7 | 100 | no |
| YBR218C | PYC2 | 1181 | MAC% | Q373K | SB | 1.7 | 78.7 | no |
|  |  |  |  | E722K | SB | 0.1 | 0 | no |
| YLL041C | SDH2 | 267 | MAC%. kinetics | K158E | SB | 1.3 | 100 | no |
| YGR014W | MSB2 | 1306 | Glycerol | S529F | SB | 1.7 | 98.9 | yes |
| YKL029C | MAE1 | 669 | MAC% | I605V | GN | 64.9 | 0 | no |

### SB proteome reveals peculiar metabolic regulations functionally connected with some causative genes

Flor yeasts are able to grow on the wine surface at the end of the alcoholic fermentation. By creating biofilm rafts, they are able to resist to high ethanol content in harsh conditions (Legras et al. 2016). For ensuring their development, they activate particular metabolic pathways (active neoglucogenesis and respiration metabolism) that are the opposite of those developed by wine yeasts during the alcoholic fermentation. Such metabolic differences have been previously reported at the metabolomic and the proteomic levels (Moreno-García, García-Martínez, Moreno, et al. 2015; Moreno-García, García-Martínez, Millán, et al. 2015; Alexandre 2013; David-Vaizant and Alexandre 2018). In order to have a broad overview of the metabolic peculiarities of the SB strain, we reanalyzed a proteomic dataset previously generated in our laboratory (Albertin, Marullo, *et al*., 2013; Blein-Nicolas *et al*., 2013, 2015). Data explored were obtained by quantifying the proteome of 25 *S. cerevisiae* strains, including SB and GN, during the fermentation of a sauvignon blanc grape juice by a shotgun proteomics approach. Samples were collected at mid-point in triplicate allowing the quantification of 1110 proteins commonly expressed (Table S7). A global Principal Component Analysis (PCA) demonstrates that SB is strongly discriminated by the two principal axes accounting for 34 % of the total inertia suggesting an outlier protein abundance respect to 24 other strains (Fig. 5, panel A). Indeed, the Abundance Fold Change Ratio (AFCR) of SB and GN *vs* the 24 other strains were compared for each of the 1100 proteins quantified. SB displays a much distinct profile since 12.9 % of its proteome reach a 2 folds change abundance (log2_(AFCR)_ +/− 1.0) while only 2.9 % of GN proteins reach this threshold (Fig S6). Thus, proteome variance of SB and GN are 0.504 vs 0.143, respectively (variance F test, pvalue <1.10^-16^). This analysis demonstrated that SB has a particular proteome compared to GN and even to other *S. cerevisiae* strains.

**Figure 5.**
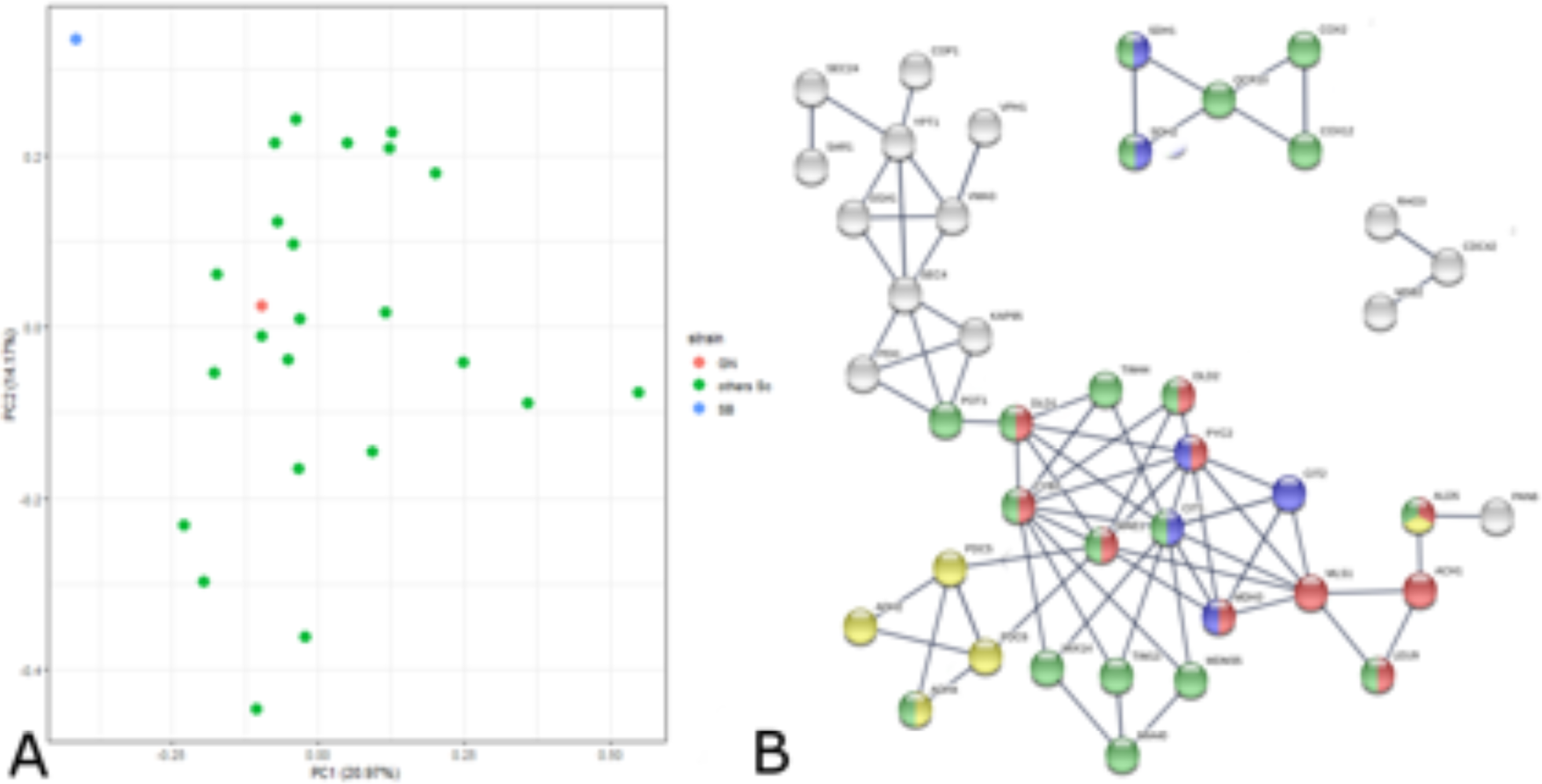
Proteomic analysis reveals the outlier behavior of the SB strain. **Panel A.** We reanalyzed a proteomic dataset previously obtained by shotgun quantitative proteomics (Blein *et al*. 2015). Yeast samples of 25 *S. cerevisiae* strains including SB and GN) were collected at mid fermentation of a Sauvignon blanc grape juice. A set of 1110 proteins common to all the strain was selected for analyzing strain relationships by a principal component analysis. The first two components representing 34% of the total inertia illustrate that the proteome of the strain SB (blue point) is quite divergent from the other *S. cerevisae* strains including GN (red point). **Panel B** The functional interactions between 207 differentially expressed proteins and the eight QTG validated in this study was interrogated by using STRING algorithm. The three clusters encompassed 2, 4 and 31 proteins showing a strong functional interaction with the four causative genes *PYC2*, *MAE1, MSB2* and *SDH2*, (black crosses). Active interactions were computed using the STRING algorithm on the base of experimental data and annotated database with a minimal interaction score of 0.8. Proteins were colored according to their mitochondrial origin (red), their involvement in pyruvate metabolism (blue) or in neo glucogenesis (green).

In order to analyze the origin of this discrepancy, we deeply compared SB and GN using the 1264 proteins quantified in both strains (Table S7). This comparative analysis reveals a set of 207 proteins with an ACFR higher than 2 (Table S8). Within this set, a significative enrichment was found for mitochondrial proteins which represent 33% of the pool (χ^2^ test=2.10^-5^). We sought functional interactions between the eight causative genes identified and the set of 207 differentially expressed proteins by performing a STRING analysis (Szklarczyk et al. 2019) (see methods). Three of the six interaction networks computed clearly linked four QTG with proteins differentially expressed (Fig 5, panel B). The main cluster, linked to the causative genes *PYC2* and *MAE1,* encompassed 31 proteins including many enzymes related to pyruvate and citrate metabolism (Mls1p, Leu9p, Ach1p, Mdh3p, Dld1p, Dld2p, Ald5p, Cyb2p, Cit1p, Cit2p). The fold change abundance of such proteins suggests the existence of differential metabolic regulations between SB and GN. For instance, three of the four *S. cerevisiae* enzymes (Dld1p, Dld2p and Cyb2p) involved in the lactate metabolism are at least 2.5 less abundant in SB. These proteins are supposed to be repressed by glucose and anaerobiosis and participate to the oxidation of lactate into pyruvate (Bekker-Kettern, 2016). Other proteins, belonging to the glyoxylate shunt and TCA, were differentially quantified (2-fold change ratio). Interestingly, the oxidative branch of TCA and the glyoxylate shunt (i.e. Mls1p, Dal7p, Cit1p, Cit2p, Aco2p) are broadly more abundant in SB while proteins participating to the reductive branch of TCA (i.e. Fum1p, Mdh1, Sdh2p) are more abundant in GN (Fig S7, panel A). These metabolic pathways are directly connected with two causative genes identified in this study *MAE1* and *PYC2* that controls *MAC%*. Strikingly, the cytosolic malate synthase Mls1p catalyzing the condensation of glyoxylate and acetyl CoA in L-malate is 7 folds more abundant in SB (log_2_(AFCR)>2.8) and would directly enhance its cytosolic pool of malic acid. These noteworthy variations of proteins abundance are not due to a singular contrast between SB and GN proteomes but reflect a clear specificity of SB central metabolism regulation. Indeed, the AFCR computed between SB and the 24 other *S cerevisiae* strains (average value) is very similar to the AFCR of SB *vs* GN (Pearson cor. test <10^-13^) (Fig S7 panel B). This analysis suggests that the peculiar proteome of SB would be due to its unusual mosaic origin encompassing large stretches of flor yeast genome.

## Discussion

### The flor yeast origin of the parental strain SB is likely involved in the diversity of carbon catabolism in the SBxGN progeny

This work aimed to identify natural genetic variations that possibly modulate the catabolism of carbon sources during wine fermentation. From an applied point of view, this goal is particularly relevant for wine industry in order to cope with two main negative effects of global warming: (*i*) the rise of ethanol content and *(ii*) the reduction of the total acidity of wines. This general trend is due to the increasing concentration of sugars coupled with a drop of malic acid content in grape juices around the world (van Leeuwen and Darriet 2016). By applying a QTL mapping strategy, eight Quantitative Trait Genes (QTG) impacting the carbon balance during the wine fermentation were identified. Although, reciprocal hemizygosity assay fails to identify candidate genes that significantly decrease the final ethanol content of wine, this study allows the identification of natural allelic variations controlling two remarkable phenotypes: the glycerol production and the percentage of malic acid consumed (*MAC%*). The schematic relationships of their respective proteins in the yeast metabolism map are shown on Fig 6.

**Figure 6.**
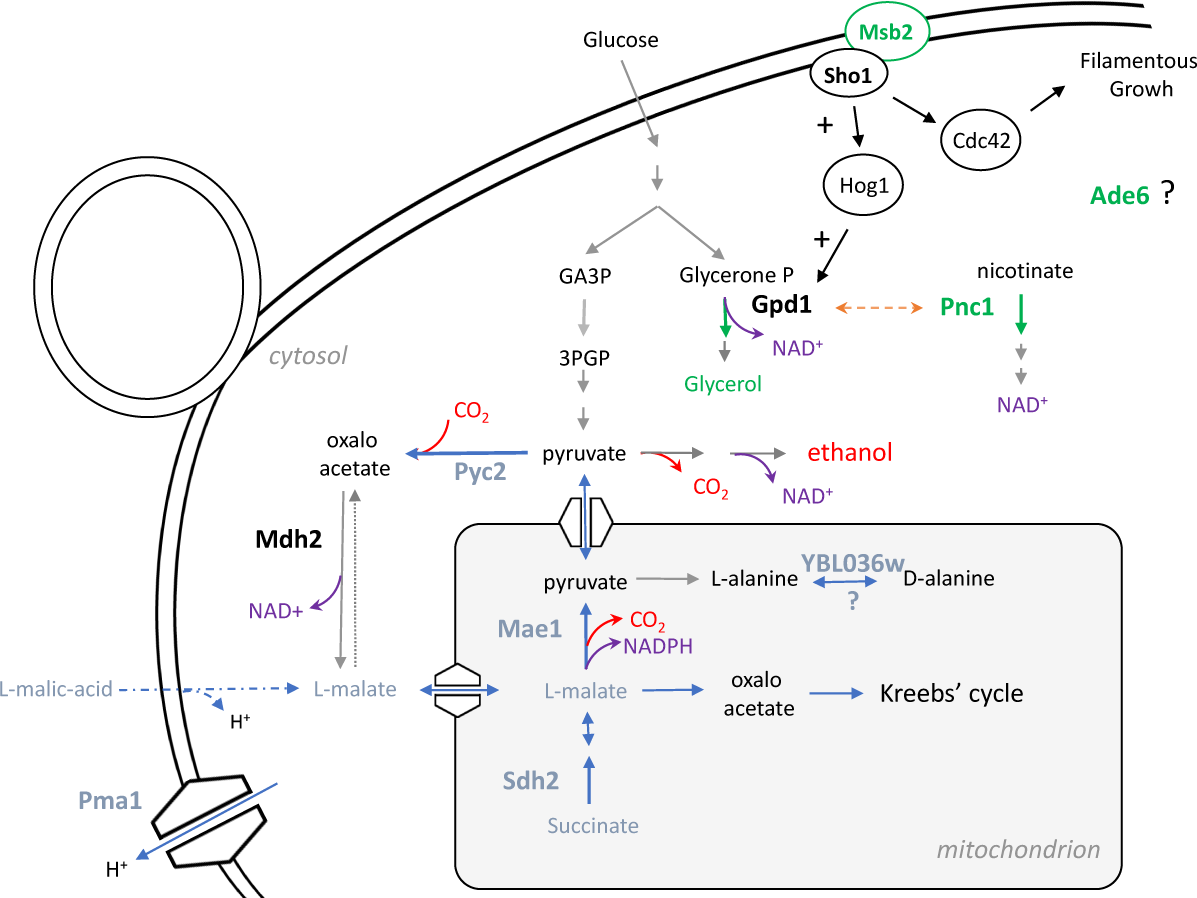
Relative position of the eight QTG in the metabolic map of *S cerevisiae*. The metabolic relationships between the eight causative genes identified in this study is presented. Genes impacting glycerol production are represented in green while genes impacting *MAC%* are presented in blue.

This study was carried out using two meiotic segregants (SB and GN) derived from commercial starters widely used in wine industry (Actiflore BO213 and Zymaflore VL1, Laffort, France). Such commercial starters have been selected in the past for their technological properties by sampling spontaneous wine fermentations (P Marullo, pers com). Unexpectedly, we find out that the SB genome has a mosaic structure inherited from two distinct groups of *S. cerevisiae* population: the wine and the flor yeasts (Peter et al. 2018). Around 40 % of the SB genome is flor specific suggesting that BO213, the parental strain of SB, would be an F1-hybrid resulting from the cross of a flor yeast and a wine yeast, as previously observed for others wine commercial strains related to the Champagne group (Coi et al. 2017).

Wine yeasts are adapted to a fast development on grape must in competition with numerous other species in a sugar rich environment and many natural allelic variations related to their adaptation to grape juice have been described in the past (Peltier et al. 2019). In contrast, flor yeasts are adapted to survive in wine, a sugar-depleted environment containing high ethanol degree and low oxygen. Thus, flor yeasts would have accumulated specific genetic variations for coping with this harsh environment. Many efforts have been made for identifying such adaptation signatures especially concerning the development of the flor velum. This biofilm-like growth is essential for reaching the wine surface and to get oxygen which is mandatory for catabolizing ethanol and producing energy (Legras et al. 2016). Allelic variations specific to flor yeasts have been detected by using comparative genomics and the role of two genes (*SFL1* and *RGA2*) participating in the regulation of *FLO11* has been demonstrated (Coi et al. 2017). In the SBxGN cross, wine and flor specific alleles segregate providing the opportunity to study the phenotypic impact of gene pools that have undergone parallel evolutionary routes with different selective pressures. Indeed, nine of the fourteen QTL identified are located in flor specific regions allowing the molecular validation of six genes (*PMA1*, *PNC1*, *PYC2*, *SDH2, MAE1,* and *MSB2*) characterized by flor specific alleles. This suggest that part of the allelic variations involved in the adaptive divergence between wine and flor yeast had been captured.

Functionally, these genes are involved in key pathways discriminating flor yeast and wine yeast metabolisms. First, *MSB2* encodes a signaling mucin protein acting as a stress or nutrient deprivation receptor (Cullen and Sprague 2012). Msb2p is associated with the transmembrane osmosensor Sho1p and transmits the signal to the downstream components of the monomeric G-proteins Rho involved in both filamentous growth (FG) and the high osmolarity glycerol (HOG) pathways (Tatebayashi et al. 2007). HOG pathway plays a key role for adaptation against high osmolarity levels by increasing the production of glycerol (Hohmann, 2009), the second more abundant metabolite of fermenting yeast after ethanol. The comparative analysis of *MSB2* sequence reveals a unique ns-SNP between the parental strains (Table 1). The SB allele *S529F* is specific to flor yeasts and lowers the glycerol production respect to the GN allele. The *MSB2^S529F^* allele has a predicted deleterious effect that would impact the signal transduction of both HOG and FG MAPK pathways. Such pathways share common components but are induced by different stimuli and provides specific responses (Pitoniak et al. 2009). The essential Rho protein Cdc42p has been described to stimulate glycerol production by triggering the MAPK Hog1p (Hohmann, 2009). Cdc42p is threefold less abundant in SB which is consistent with the hypothesis of a low Msb2p activity in this background. In contrast the non-essential GTPase Rho3p also involved in cell polarity is three times more abundant in SB. Interestingly, the abundance fold ratio of Rho3p and Cdc42p are specific to SB (compared to others *S cerevisiae* strains) and might be related to the filamentous growth specificities of flor yeast required for the velum formation.

A flor-specific allele was also found in the sequence of *PNC1* which encodes for a nicotinamidase that converts nicotinamide to nicotinic acid. Pnc1p, which is induced by the osmotic stress, restores redox balance by regenerating NAD^+^ from nicotinamide via the NAD^+^ salvage pathway (Effelsberg et al. 2015; Ghislain, Talla, and François 2002). RHA reveals that the allele *PNC1^SB^* enhances both the glycerol production and the *MAC%*. A direct functional link exists between *PNC1* and glycerol biosynthesis since this protein is co-imported in the peroxisome with Gpd1p, a major controlling enzyme of glycerol biosynthesis (Nevoigt and Stahl 1997). Under osmotic stress, their overexpression saturates the peroxisome importation system and therefore this protein became cytosolic and active (Effelsberg *et al*., 2015). The role of Pnc1p in *MAC%* is more complex to explain and might be linked to the NAD^+^/NADH^+^ homeostasis itself that is tightly controlled (Bakker et al. 2001). This organic acid can be oxidized in pyruvate (by the malic enzyme Mae1p) or in oxaloacetate (by malate dehydrogenases). Thus, an active malic acid consumption would increase the intracellular levels of NADH^+^ requiring an increase of glycerol production for regenerating the NAD^+^ pool.

Another flor yeast specific allele impacting *MAC%* is *MAE1* that encodes for the mitochondrial malic enzyme that catalyzes the oxidative decarboxylation of malate to pyruvate (Boles, de Jong-Gubbels and Pronk, 1998) achieving the malo-ethanolic fermentation (Volschenk, Vuuren and Viljoen–Bloom, 2003). Interestingly, *MAE1* was also reported to influence the formation of higher alcohols, fusel acids, and acetate esters in another mapping population where the same SNP is segregating (MAE1^I605V^) (Eder *et al*., 2018). These data suggest that this allelic variation would have pleiotropic consequences in an enological context, affecting the malic acid consumption as well as the biosynthesis of relevant wine volatile compounds.

A second pleiotropic gene to be discussed is *PMA1* which encodes for a membrane ATPase the major regulator of cytoplasmic pH and plasma membrane potential. During wine fermentation, pH has a great impact on intracellular malic acid diffusion and consumption (Salmon, 1987; Delcourt *et al*., 1995; Saayman and Viljoen-Bloom, 2006). Indeed, malic acid charge is strongly dependent of the wine pH since the *pka*1 of this diacid is 3.54. Bellow a pH value of 3.4, the entry of a malic acid molecule in the cytoplasm result to a net proton influx that must be pumped over for maintaining pH homoeostasis with an energy cost of 1 ATP per molecule. In the present work, the QTL VII_482 related to *PMA1* has the strongest effect observed with a positive impact of the GN allele on malic acid consumption. Previously, we demonstrated that *PMA1* inheritance influences fermentation kinetics with a strong interaction with the pH of the medium. Indeed the GN and SB alleles increase the fermentation rate when the pH are 3.3 and 2.8, respectively (Martí-Raga *et al*., 2017). These fine grain gene-environment interactions might result from the consumption level of malic acid in relation with the pH of wine.

Two other genes with a direct connection with malic acid metabolism were shed on light. The gene *PYC2* involved in gluconeogenesis pathway encodes for a pyruvate carboxylase that converts pyruvate to oxaloacetate (Stucka *et al*., 1991; Walker *et al*., 1991). During fermentation, pyruvate carboxylase is the sole source of oxaloacetate playing an essential role in aspartate biosynthesis, TCA turnover, and malic acid biosynthesis (Huet *et al*., 2000). Indeed, *PYC2* overexpression enhances malic acid production in a bioengineering context (Bauer *et al*., 1999). We hypothesized that the allelic variants of SB may have reduced the Pyc2p activity reducing the biosynthetic flux of malic acid from pyruvate. To cope with this reduction, a first metabolic alternative would be the *de novo* synthesis of malic acid from the glyoxylate shunt. This is consistent with the high abundance of the malate synthase (more than 7 folds) observed in SB respect to GN. A second metabolic alternative would be a strongest uptake from the external media which is the hallmark of the SB strain.

Finally, a surprising effect of *SDH2* deletion was observed. This gene encodes for a subunit of the succinate dehydrogenase complex (complex II) ensuring electron transfer from succinate to ubiquinone. This TCA cycle step is involved in the mitochondrial respiratory chain and is mostly inactive during the alcoholic fermentation (Camarasa, Grivet and Dequin, 2003) due to oxygen depletion and catabolic repression (Klein, Olsson and Nielsen, 1998; Kwast, Burke and Poyton, 1998). Indeed, under sake brewing conditions, the CO_2_ production rate was not impacted in double mutants *Δsdh1, Δsdh2* (Kubo, Takagi and Nakamori, 2000). These commonly admitted results contrasted with the strong haploinsufficiency effect of *SDH2* deletion observed for *MAC%* and fermentation kinetics in M15 medium (Fig S5). Although we could not measure a significant difference between hemizygous hybrids, the strong haploinsufficiency observed suggests that the succinate dehydrogenase complex would play an unsuspected physiological role in this specific background. Interestingly STRING analysis reveals that five proteins functionally associated to *SDH2* are differentially synthetized between SB and GN. These proteins belong to the respiratory complexes II, III and IV. Thus, complex II (Sdh1p and Sdh2p) is less abundant in SB while proteins belonging to complex III (Qcr10p) and IV (Cox2p and Cox12p) are more abundant. Due to the functional importance of protein stoichiometry in such complexes, abundance change in few proteins would impact the residual activity of the respiratory chain. Therefore, the functional understanding of the succinate dehydrogenase complex during alcoholic fermentation will require further analyses that are not the purpose on the present paper.

Flor yeasts exhibit an active gluconeogenesis and respiration catabolism during velum development that impact their proteomics response (Legras et al. 2016; Alexandre 2013) (Moreno-García, García-Martínez, Moreno, et al. 2015). However, to our knowledge, a comparative proteomic study between flor and wine yeast was never achieved. Since the SB strain harbor 40% of the genomic signature of a flor yeast, we supposed that this strain could exhibit particular flor yeast features at the proteomic level. This prompted us to compare the fermentation proteome of SB with other *S cerevisiae* strains including the parental strain GN used in this study. A large comparative proteomics study between strains of the same species carried out in our laboratory was reanalyzed for this purpose (Blein et al. 2015). The abundance of 1100 proteins commonly quantified in 25 *S. cerevisiae* strains clearly demonstrated that SB exhibit a peculiar proteomic regulation (Fig 5, panel A) during wine fermentation (Table S7). Strikingly most of the proteins differentially regulated between SB and GN are due to the specific proteomic patterns of SB discarding the fact that the SB vs GN proteomic variations would be due to the GN strain (Fig S6, panel B). Several proteins involved in pyruvate and gluconeogenesis were differentially quantified. Many of them have been previously described as specific signature of velum development (Moreno-García, García-Martínez, Moreno, et al. 2015).

By implementing a STRING analysis, we attempted to retrace a functional link between the eight QTG identified and the proteomic variations observed between parental strains. This indirect analysis would bridge the gap between specific flor yeast variations and the overall proteomic discrepancy of the SB strain. Three causative genes (*MSB2*, *SDH2* and *PYC2*) harboring flor specific alleles were functionally connected with three protein clusters (Fig 5, panel B). *PYC2* and *SDH2* are directly involved in central carbon metabolism playing an essential role in gluconeogenesis and respiration, respectively. The first controls the unique way for producing glucose from ethanol since the pyruvate kinase catalyzed an irreversible reaction (Pronk, Steensma, and Van Dijken 1996). The second belongs to the succinate dehydrogenase which is inactivated during the fermentation and that constitutes the first step of respiration chain (complex II) which is essential for producing energy in aerobic conditions. A contrasted regulation between the oxidative and reductive branch of TCA was observed in the strain SB (Fig S7A) promoting the idea that succinate dehydrogenase activity would participate to the regulation of TCA proteome. Although this hypothesis remains to be validated by further experiments, we hypothesized that the specific flor alleles Sdh2^K158E^ and Pyc2^Q373K^ carried by SB strain might impact the overall proteomic response of this strain by controlling key steps of gluconeogenesis and TCA cycle.

## Materials and Methods

### Yeast strains and culture media

All the strains used in this study belong to the yeast species *Saccharomyces cerevisiae*. SB and GN strains are monosporic clones derived from industrial wine starters, VL1 and Actiflore BO213, respectively. Generation of the SBxGN and segregant populations were described by (Peltier, Sharma, *et al*., 2018). Briefly, F1-hybrids were obtained by manual crossing with micromanipulator. After sporulation on ACK (2 % potassium-acetate, 2% agar) media, monosporic clones were isolated by micromanipulation. Yeast was cultured at 30 °c in yeast YPD media (10 g/L yeast extract, 20 g/L peptone and 20 g/L glucose) and solidified with 2 % agar when required. The strains were stored long term in YPD with 50% of glycerol at − 80 °C.

### Phenotyping

The two grape juices used, Merlot of vintage 2015 (M15) and Sauvignon Blanc of vintage 2014 (SB14), were provided by Vignobles Ducourt (Ladaux, France) and stored at − 20 ° C. Before fermentation, grape juices were sterilized by membrane filtration (cellulose acetate 0.45 μm Sartorius Stedim Biotech, Aubagne, France). Fermentations were carried out as previously described (Peltier, Bernard, et al. 2018). Briefly, fermentations were run at 24 °C in 10 mL screw vials (Fisher Scientific, Hampton, New Hampshire, USA) with 5 mL of grape must. Hypodermic needles (G 26–0.45 × 13 mm, Terumo, Shibuya, Tokyo, Japan) were inserted through the septum for CO2 release. Two micro-oxygenation conditions were used by applying or not constant orbital shaking at 175 rpm during the overall fermentation. For this data, three fermentation conditions were used: SB14 with shaking (SB14_Sk), M15 with shaking (M15_Sk) and M15 without shaking (M15). Fermentation progress was estimated by regularly monitoring the weight loss caused by CO2 release using a precision balance. The amount of CO2 released over time was modeled by local polynomial regression fitting with the R-loess function setting the span parameter to 0.45. From this model *CO2max* parameter was extracted: maximal amount of CO2 released (g.L^-1^) and the end of the fermentation. Fermentation conditions were described by (Peltier, Sharma, et al. 2018). Glycerol and malic acid concentration were determined by enzymatic assay (Peltier et al. 2018) using K-GCROLGK and K-LMAL-116A enzymatic kits (Megazyme, Bray, Ireland), following the instructions of the manufacturer.

### Linkage analysis

The QTL mapping analysis was performed with the R/qtl package (Broman et al. 2003) on the data collected in the three environmental conditions by using the Haley-Knott regression model that provides a fast approximation of standard interval mapping (Haley and Knott 1992). The analysis is taking in account environment and cross as an additive covariate, aiming to identify QTL robust to environment and cross factor:

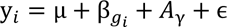

Where y_*i*_ is the phenotype for individual *i*, µ the average value, β*_!_* the QTL genotype for individual ਍, *A*_#$_ the matrix of environment covariates (y = M15_Sk, SB14_Sk, M15) and ɛ the residual error. For each phenotype, a permutation test of 1000 permutations tested the significance of the LOD score obtained, and a 5% FDR threshold was fixed for determining the presence of QTLs (Churchill and Doerge 1994). The QTL position was estimated as the marker position with the highest LOD score among all markers above the threshold in a 30 kb window.

### Hemizygous hybrids construction

For each QTL, candidate genes were sought in a 30 kb windows around the QTL position with the maximal LOD score. Genes with non-synonymous SNPs and/or with a function related to the trait of interest were retained. Candidate genes were validated by reciprocal hemizygosity analysis according to (Steinmetz et al., 2002) using SBxGN hybrid. Deletion cassettes were obtained by PCR amplification of the disruption cassette plus 500 pb of the flanking regions using as genomic template the genomic DNA of the strains Y04691, Y03717, Y04878, Y03751, Y04405, Y01529, Y03062 of the EUROSCARF collection (http://euroscarf.de), which contain disruption cassettes for the following genes: *ADE6*, *GPM2*, *MAE1*, *MCH1*, *PNC1*, *PYC2*, *SDH2*, *YBL036C*, respectively. Primers used for strains construction are listed in File S2. Reciprocal hemizygotes for *MSB2*, *PDR1* and *PMA1* were previously constructed with the same strategy by (Martí-Raga *et al*., 2017).

### Phylogenic analysis

Publicly available sequences of yeasts from wine and flor genetic groups were retrieved from (Peter et al. 2018; Legras et al. 2018) and are listed in table S6. A matrix of 385,678 SNPs was generated with GenotypeGVCFs from GATK after gvcf files were constructed as detailed in (Peter et al. 2018). This matrix was used to build a neighbor-joining tree using the *ape* and *SNPrelate* R packages. Flor and wine yeast genetic groups were determined according to (Peter et al. 2018; Legras et al. 2018) and correspond to the flor genetic group and the Wine/European (subclade 4), respectively. Flor yeast specific alleles were defined as alleles with a frequency difference of 90 % between flor and wine genetic groups.

### Proteomic data reanalysis

The dataset used for reanalyzing proteome specificities of the strain SB correspond to the supplementary *table S5* published by Blein *et al*. (2015). This dataset compassed the proteomes of 66 *Saccharomyces* strains quantified during the alcoholic fermentation of a Sauvignon blanc grape juice at two temperatures. Among those strains, 28 *S cerevisiae* strains constituting a half-diallel design of 7 parental strains of different origins and 21 F1-hybrids. In that study the parental strains SB and GN were referenced as E2 and E3, respectively. A subset portion of this large data set was reanalyzed for narrowing down the proteomic specificities of the strain SB. Only the proteome corresponding to *S cerevisiae* strains measure at 26°C were kept. Indeed, proteomic data for the strain E2 (SB) at 18°C were not available. In addition, we removed the proteomes of the strains W1, EW21 and EW31 due to the lower number of proteins quantified (<900) respect to the other strains. By applying these filters, we analyzed the abundance of 1100 proteins commonly quantified in 25 *S. cerevisiae* strains including GN and SB. In addition, the list of the 1264 proteins specifically detected between SB and GN was listed in the table S7. The abundance values indicated in are the average of three biological replicates where 90% of the data points have a CV% lower than 5.37. The Abundance Fold Change Ratio (AFCR) of the strains SB and GN were expressed in log2 for an easier comparison. An arbitrary AFCR threshold of +/-1 was used for selected proteins having a relevant abundance change, this basic threshold is widely used in the proteomics literature. The table S8 provides the list of the 207 proteins selected in the set of the 1264 proteins common to SB and GN. Proteins with a differential abundance between SB and GN were used for computing a STRING analysis in order to find out functional connections with the eight genes validated in this study. The permanent link of such analysis is the following https://version-11-0.string-db.org/cgi/network.pl?networkId=pEeVlh8dPgJJ. The interaction classes interrogated were “experiments” and “databases” with the highest confidence score.

### Statistical analyses

All the statistical and graphical analyses were carried out using R software (R Core Team 2018). The *lato sensu* heritability *h^2^* was estimated for each phenotype as follows:

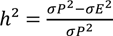

where *σP^&^* is the variance of progeny population in each environmental conditions, explaining both the genetic and environmental variance of the phenotype measured, whereas *σE^2^* is the median of the variance of replicates in each environmental conditions, explaining only the environmental fraction of phenotypic variance.

## Acknowledgements and funding information

The authors thank Justine Pape, Dylan Dos Reis and Elodie Kaminski that helped managing fermentations. This work was funded by Région d’Aquitaine (https://www.nouvelle-aquitaine.fr). The funders had no role in study design, data collection and analysis, decision to publish, or preparation of the manuscript.

a ns-SNP have been predicted to be to have a deleterious effect on protein according to PROVEAN algorithm

## Supplementary Figure

**Fig S1. Correlation between traits.**

Correlation between traits. Data is normalized according to environment. Each dot represents the average value of an individual in one of the three phenotypic condition. Correlation coefficient and P value of Spearman’s correlation test is indicated. *CO2max* is negatively correlated with *glycerol* and positively correlated with *MAC%* (Spearman test, pval < 0.01). However, *rho* values observed are quite low (<0.2) because the variation in CO2 production is balanced by glycerol production and malic acid consumption.

**Fig S2. Distribution of traits.**

**Left**. Distribution of the progeny according to trait and media is represented. Dashed vertical line represent parental average value. **Right**. Data is normalized according to environment. Distribution of the progeny in all media, according to trait and cross. Dashed vertical line represent parental average value.

**Fig S3. QTL effect in population.**

Effect of each QTL according to parental inheritance. Each dot represents the phenotypic value of one individual and are colored according to their marker inheritance. Bigger points represent the mean of the population.

**Fig S4. Discrepancy for MSB2**

**Panel A**. Effect of the marker associated to *MSB2* in the offspring. Each dot represents the phenotypic value of one individual and are colored according to their marker inheritance. **Panel B.** Result of RHA test for *MSB2*. The represented value is from at least 5 biological replicates. The level of significance is indicated as follows: * *p* ≤ 0.1. ** *p* ≤ 0.05. *** *p* ≤ 0.01. Solid lines of kinetic curves represent the mean and the shadow the standard error.

**Fig S5. *SDH2* hemizygotes show a substantial haploinsufficiency according to media.**

The represented value is from at least 5 biological replicates. A Wilcoxon–Mann–Whitney test was applied to assess the significance of the phenotypic difference between wild type and hemizygote. The level of significance is indicated as follows: * *p* ≤ 0.1. ** *p* ≤ 0.05. *** *p* ≤ 0.01. Solid lines of kinetic curves represent the mean and the shadow the standard error.

**Fig S6. SB proteome exhibit a strongest variability than GN respect to 24 others S cerevisiae proteomes.**

The plot represents the distribution of the Abundance Fold Change Ratio (expressed in log2) of the strains SB and GN respect to the average values of 24 other strains. The variance of SB and GN computed for the 1110 proteins indicated a highest variability of the SB proteome (F-test analysis <1.10^-7^).

**Fig S7. Abundance of proteins belonging to the oxidative and reductive branches of TCA in SB respect to GN and others S cerevisiae strains**

Panel A. Abundance fold ratio of quantified proteins belonging to the TCA and the glyoxylate shunt; red and green colors indicated over and under expressed proteins in the SB strain *vs* GN (left box) or *vs* the average value of 24 *S cerevisiae* strains (right box). Panel B. correlation between the AFCR (log2) of SB *vs* GN and SB *vs* 24 *S. cerevisiae* strains for the commonly expressed proteins.

## Supplementary file

**File S1. Assessment of the alcoholic fermentation yield and variability of carbon use in wine fermentation**

**File S2. Hemizygotes construction**

## Supplementary tables

**Table S1. Genotype data of offspring**

**Table S2. Phenotype data of offspring**

**Table S3. Heritability**

**Table S4. QTL list**

**Table S5. Candidate genes**

**Table S6. Strains used for phylogeny analysis**

**Table S7. Protein dataset**

**Table S8. Protein difference SB vs GN**

## Data Availability Statement

**Phylogenic analysis**

