## Supplementary figures and images for "Flor yeasts rewire the central carbon metabolism during wine alcoholic fermentation"

### Fig S1

Correlation coefficient = −0.2 (pval = 0 )

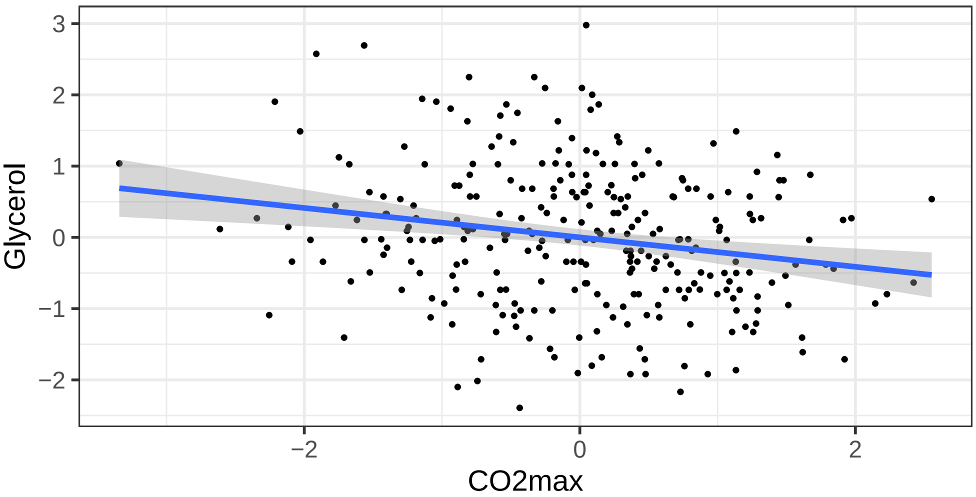


Correlation coefficient = 0.25 (pval = 0 )

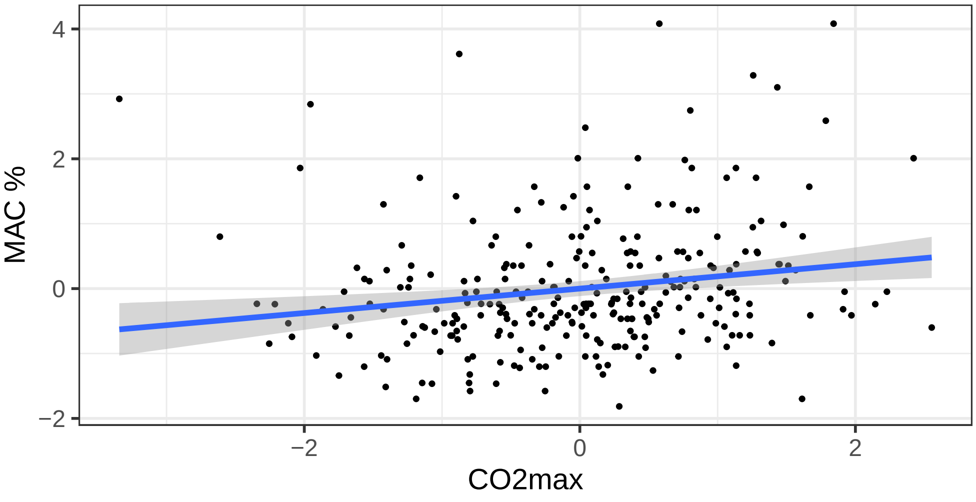

### Fig S2

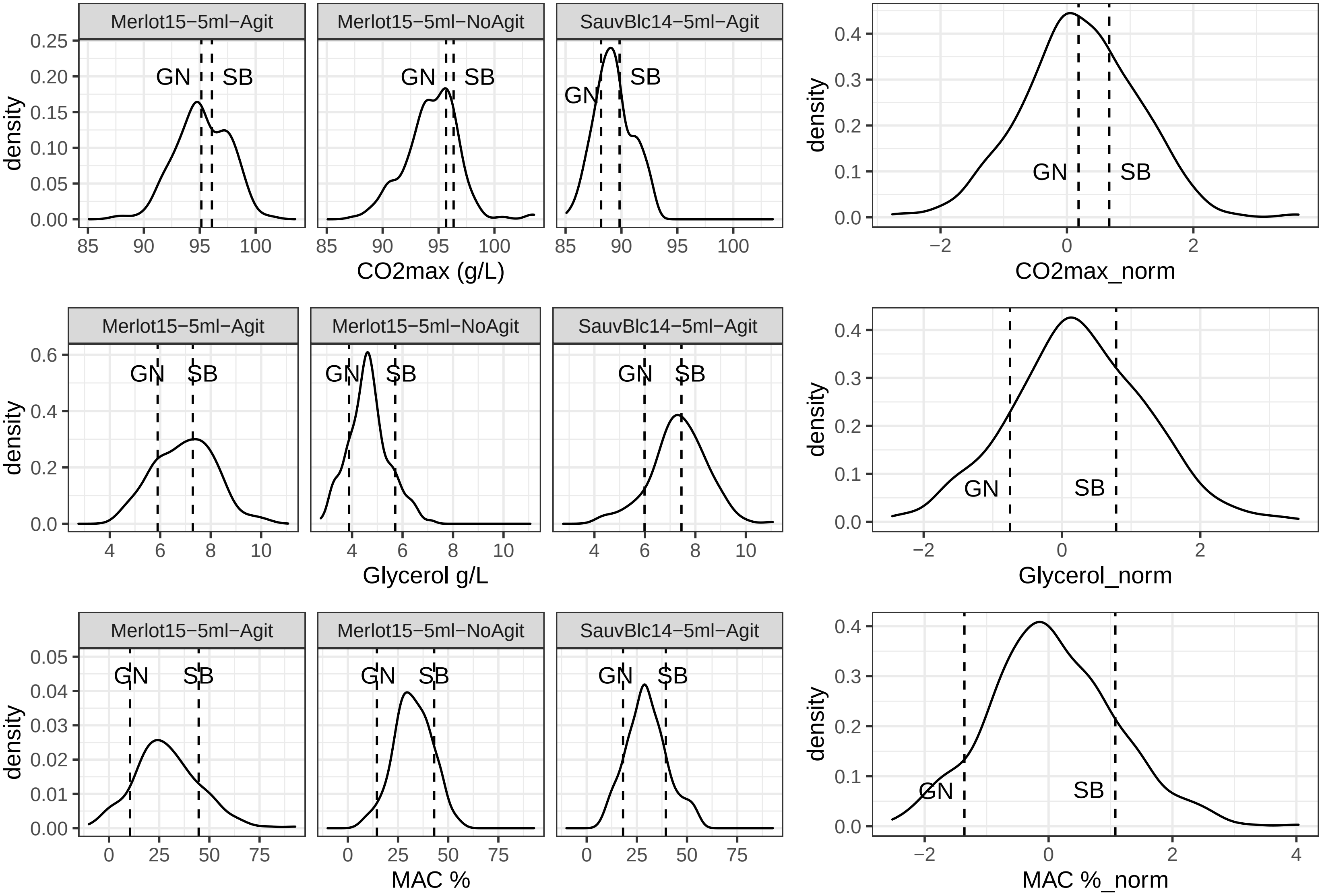

### Fig S3

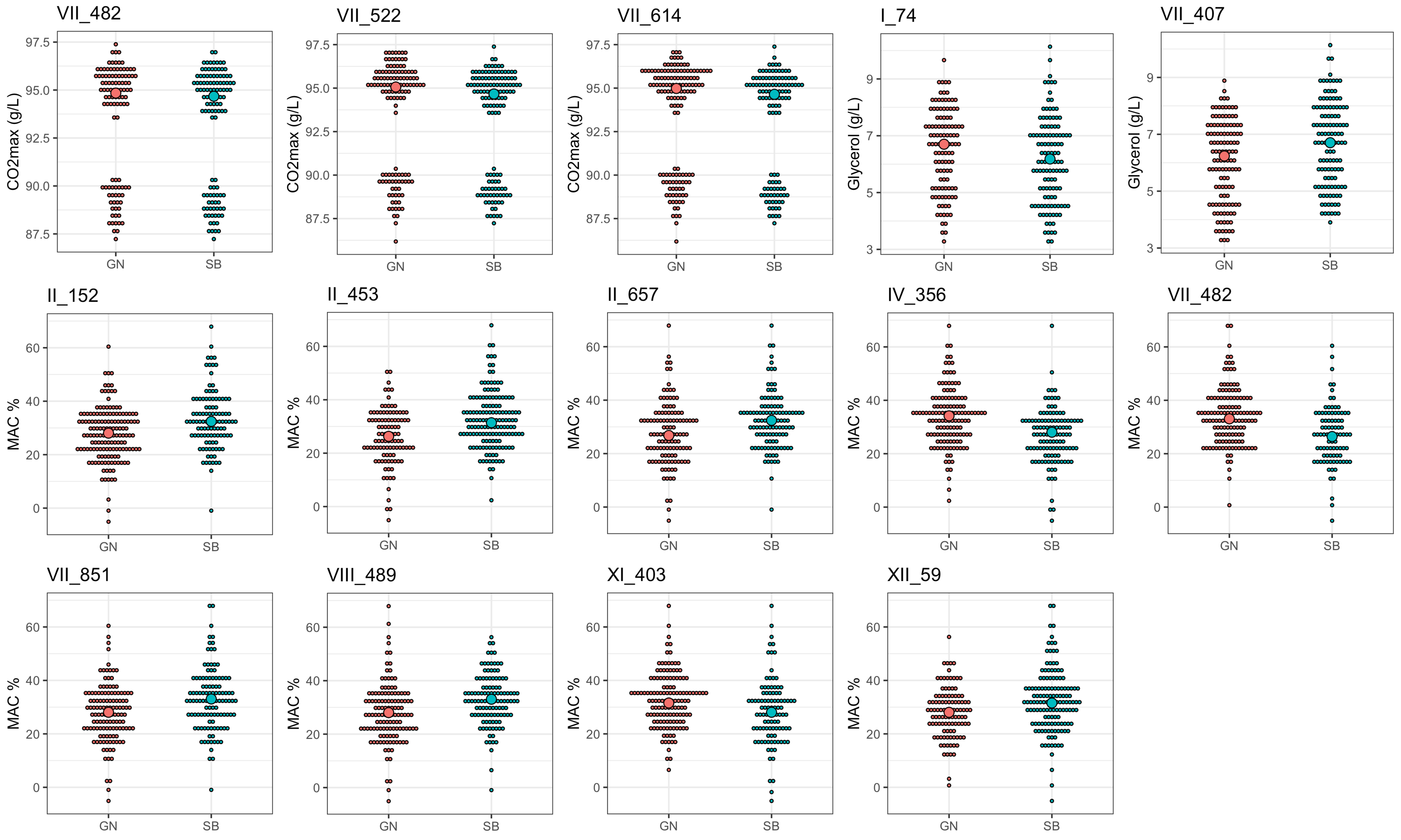

### Fig S4

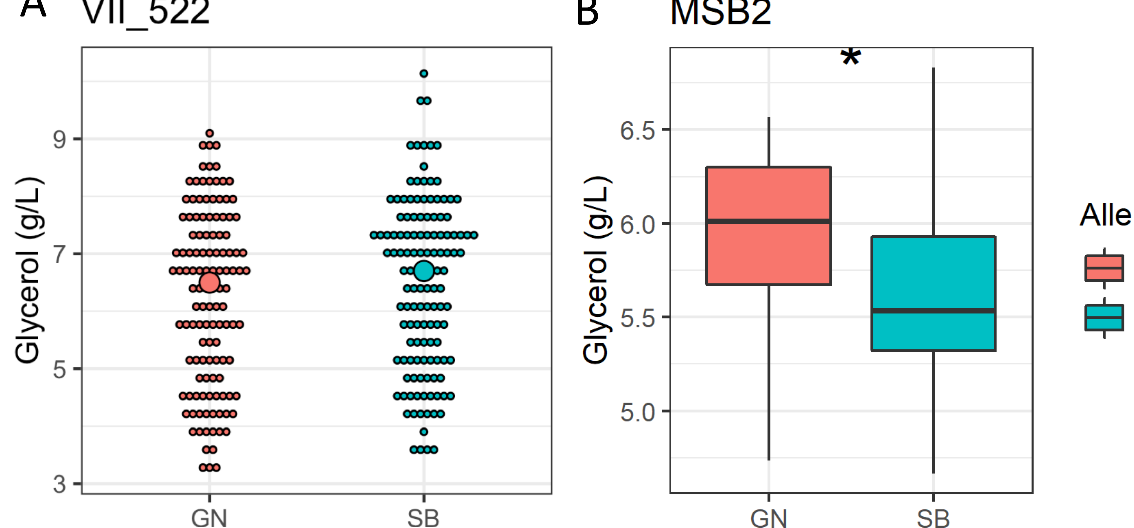

### Fig S5

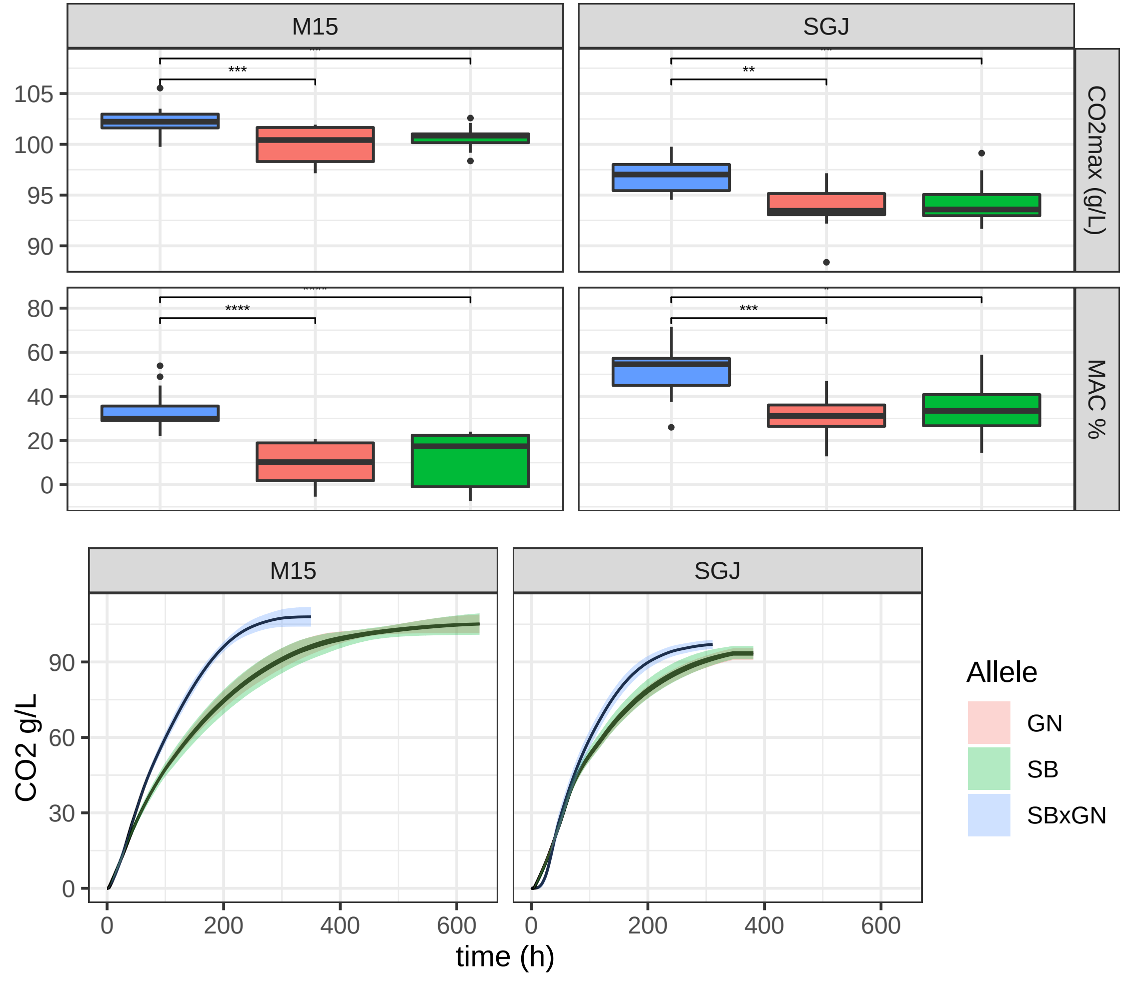

### Fig S6

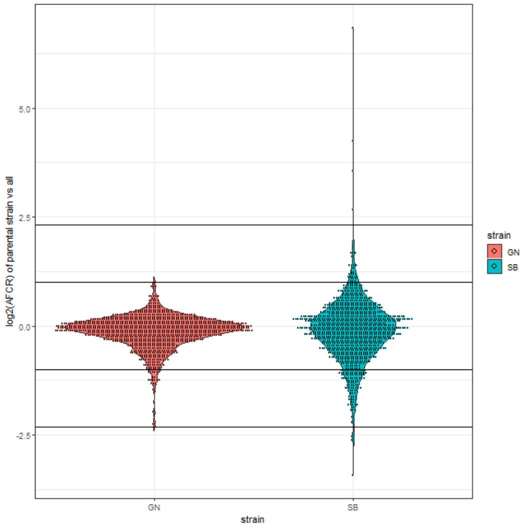

### Fig S7

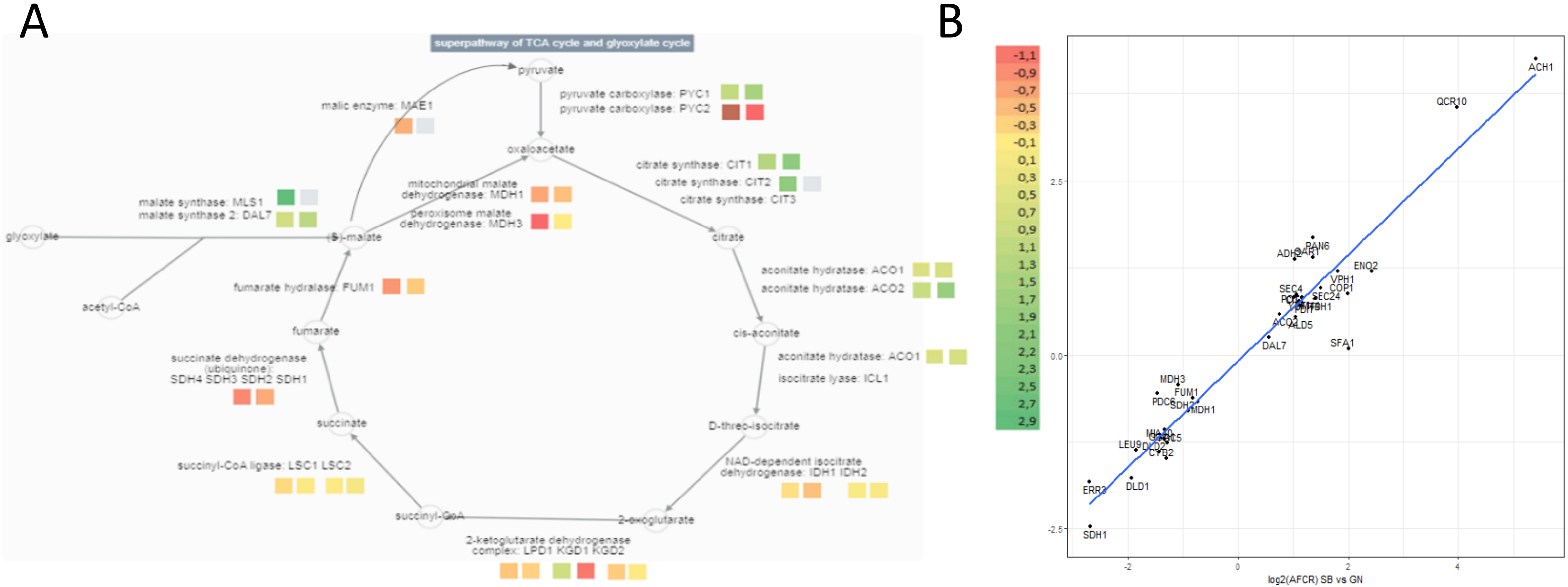
