## Supplementary material for "Flor yeasts rewire the central carbon metabolism during wine alcoholic fermentation": File S1

### Assessment of the alcoholic fermentation yield and variability of carbon use in wine fermentation

The main organic compounds assimilated and/or produced by yeast using semi-automated enzymatic assays as described in Peltier et al. (2018). The values measured for each *strain x media* combination are provided in (Table S2). For each fermentation, the concentration of residual sugars (glucose and fructose) at the end of the fermentation was lower than 2 g/L and all of them were considered as complete from an enological point of view.

A two-way analysis of variance was applied for estimating the effect of environmental (*media*) and genetic (*strain*) factors influencing each trait (see below). Since the carbon input is mostly related to glucose and fructose, grape juice has a massive effect (99% of variance explained) due to the difference of sugar concentration between M15 and SB14 grape juices. However, yeast strain significantly modulated the amount of carbon assimilated (0.4% of total variance observed). This modulation would be due to the quantity of malic acid assimilated that strongly varies according to the strain (6% of total variance observed). This organic acid is converted into ethanol through the malo-ethanolic fermentation [2]. In some cases, the malic acid consumed represented up to 1.6% of the total carbon input and was then considered as a minor but significative source of carbon.

| | Trait | M15 | | M15_Sk | | SB14_Sk | | Condition effect (%) | Strain effect (%) |
|---|---|---|---|---|---|---|---|---|---|
| | | mean | CV | mean | CV | mean | CV | | |
| Carbon in | Malic acid consumed (g/L) | 0.76 | 25.12 | 0.68 | 48.66 | 2.57 | 16.13 | 87 | 6 |
| | Glucose consumed (g/L) | 109.5 | 0 | 109.5 | 0 | 97 | 0 | 100 | 0 |
| | Fructose consumed (g/L) | 109.5 | 0 | 109.5 | 0 | 97 | 0 | 100 | 0 |
| | Carbon input (in mole) [a] | 7.34 | 0.32 | 7.34 | 0.54 | 6.62 | 0.74 | 49.1 | 17.3 |
| Carbon out | $CO_2$ produced (g/l) | 94.4 | 1.84 | 95.21 | 1.56 | 89.42 | 1.28 | 54.6 | 16 |
| | Glycerol produced (g/L) | 4.65 | 14.66 | 6.97 | 15.27 | 7.4 | 12.81 | 51.1 | 21.5 |
| | Pyruvate produced (g/L) | 0.06 | 71.87 | 0.07 | 64.51 | 0.03 | 85.29 | 5.5 | 56.8 |
| | Acetic acid produced (g/L) | 0.27 | 32.44 | 0.17 | 47.13 | 0.16 | 38.3 | 17.7 | 44.6 |
| | Carbon output (in mole) [b] | 6.59 | 1.77 | 6.72 | 1.63 | 6.34 | 1.24 | 46.1 | 19.5 |
| | Yield Ethanol/Subtract (g/g) [c] | 0.45 | 1.85 | 0.45 | 1.53 | 0.48 | 1.23 | 49.1 | 17.3 |

[a] mole of carbon of glucose, fructose, and malic acid

[b] mole of carbon of glycerol, ethanol, and $CO_2$

[c] amount of ethanol produced is inferred from $CO_2$

The carbon output was mostly due to the production of *glycerol*, *ethanol*, *$CO_2$max*, and in a much less extend to other organic compounds such as *pyruvate* and *acetic acid.* These two last compounds contribute to carbon output in a neglectable manner (0,003 and 0,02 %, respectively). Ethanol produced was not included in the dataset because its quantification by enzymatic assay was not sufficiently accurate (CV>20%). For estimating the carbon balance, the number of moles of ethanol produced was inferred by using the $CO_2$ produced assuming that pyruvate respiration is not present during wine fermentation. In order to include the contribution of malic acid consumption, the yield of alcoholic fermentation was expressed as the quantity of ethanol produced for 1 g of subtract input (sugar + malic acid). The values of fermentation yield ranged between 0.45 and 0.47 which is close to values observed in other studies [3]. Strikingly, the variance of carbon output within each condition was much higher than carbon input. This is explained by the strong yeast strain effect observed for *$CO_2$max* and *glycerol* production (16 and 21.5 % of total variance explained, respectively).


1. Peltier E, Bernard M, Trujillo M, Prodhomme DD, Barbe J-C, Gibon Y, et al. Wine yeast phenomics: a standardized fermentation method for assessing quantitative traits of Saccharomyces cerevisiae strains in enological conditions. Schacherer J, editor. PLoS One. 2018;13: 191353. doi:10.1101/191353
2. Volschenk H, Vuuren HJJ van, Viljoen–Bloom M. Malo-ethanolic fermentation in Saccharomyces and Schizosaccharomyces. Curr Genet. 2003;43: 379–391. doi:10.1007/s00294-003-0411-6
3. Tilloy V, Ortiz-Julien A, Dequin S. Reduction of ethanol yield and improvement of glycerol formation by adaptive evolution of the wine yeast Saccharomyces cerevisiae under hyperosmotic conditions. Appl Environ Microbiol. 2014;80: 2623–32. doi:10.1128/AEM.03710-13
