## Supplementary material for "Flor yeasts rewire the central carbon metabolism during wine alcoholic fermentation": File S2

### File S1 MOLECULAR TECHNIQUES

#### 1. PCR TO AMPLIFY DELETION CASSETTES TO TRANSFORM

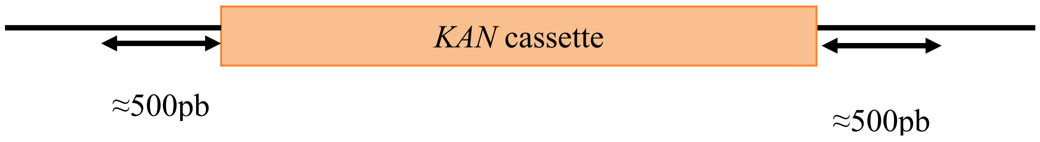


- Primers used:

Table 1. Primers used to amplify each gene specific deletion cassette.

| Gene | Sequence | Type |
|---|---|---|
| ADE6 | ATCGAGACTCGGTTGTGTCG | Forward |
| ADE6 | GGTCCAGAAGCTAAGGCCTCC | Reverse |
| GPM2 | CCAAGGCTCGACAAGGATGT | Forward |
| GPM2 | CCGTGTGGTGCCCAATTTTC | Reverse |
| MAE1 | TACTCTTCCCTAGGCGGTTT | Forward |
| MAE1 | ATCCGGACATCACACCCAAC | Reverse |
| MCH1 | TGGCAGAGTTTCAACAGCCA | Forward |
| MCH1 | TGGGATACTTGGTGAATTCCGT | Reverse |
| PNC1 | GTGGCACACAGGGTAATGAA | Forward |
| PNC1 | TGCTCTTGAAATGAAAACGGAA | Reverse |
| PYC2 | TGTCACTAACGACGTGTCCC | Forward |
| PYC2 | CCCATTGGTTCTATTGGGCAG | Reverse |
| SDH2 | ATTGCTGAGGTGCAAATGGC | Forward |
| SDH2 | TGGTGTTCTCTTCTCATTGCT | Reverse |
| YBL036C | GAGCAACAGGTAACAGGGGA | Forward |
| YBL036C | GCGATGCTTTGGGAAAAGAGG | Reverse |

#### 2. PCR TO VERIFY TRANSFORMATION

Before transformation:

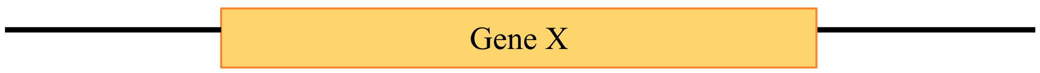

After transformation

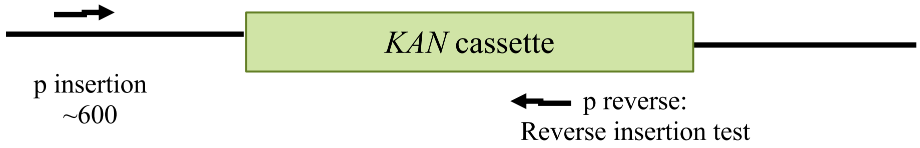


The PCR verification consist in using one primer at 600 pb (approx.) from the deleted gene loci and one primer that will anneal inside the *KAN* cassette. Thus, we verify the correct insertion of the KAN cassette in the right loci.

- Primers used:

Table 2. Primers used in the verification of the transformation.

| Gene | Sequence | Type |
|---|---|---|
| All | CGGCGCAGGAACACTG | Anneals at the middle of the *KAN* cassette. |
| ADE6 | CGACAAGAGTCCAAGAGGCAA | Insertion test |
| GPM2 | CCACTCATTCAGAGGCTGTC | Insertion test |
| MAE1 | TCGTGCATTGCAAGGTTTTT | Insertion test |
| MCH1 | TAGTTCCACCGCCAATGGAT | Insertion test |
| PNC1 | CCTCTTTCCCTACGATCCTCC | Insertion test |
| PYC2 | TGTACTACAGGAAGCAGAAACA | Insertion test |
| SDH2 | GGCTCAAGACCGTGAATGGA | Insertion test |
| YBL036C | GTTTGTTGGTGGGTTCTCGC | Insertion test |

#### 4. RFLP / qPCR

EITHER RFLP OR qPCR TO CHECK IF THE REMAINING ALLELE IN THE TRANSFORMANT IS SB OR GN.

Table 3. Primers used in the RFLP

| Gene | Sequence | Type | Enzyme | Allele cut |
|---|---|---|---|---|
| MAE1 | CGCTGCAGTGGACCAACT | RFLP forward | MboI | SB |
| MAE1 | GCTAAGCACGATGTGTATGTTTTAT | RFLP reverse | MboI | SB |
| MCH1 | TCTCAACCGTGGCAGAAACA | RFLP forward | PVUII | SB |
| MCH1 | TGGGATACTTGGTGAATTCCGT | RFLP reverse | PVUII | SB |
| PNC1 | TCTCCAGATACGATTATGATGTGCT | RFLP forward | CviKI-1 | SB |
| PNC1 | GGAGAGTGGTAGGTGTATGTTGA | RFLP reverse | CviKI-1 | SB |
| PYC2 | TGTCACTAACGACGTGTCCC | RFLP forward | MmeI | GN |
| PYC2 | GATGAGCGTCTCTCCAGGTG | RFLP reverse | MmeI | GN |

Table 4. Primers used in the qPCR

| Gene | Sequence | Type |
|---|---|---|
| ADE6 | TGACTTGGGTGCTAAATTCGATATTAGAAAGG | GN specific forward |
| ADE6 | TGACTTGGGTGCTAAATTCGATATTAGAAAGA | SB specific forward |
| ADE6 | TAACACCGACATCCGCAACA | qPCR reverse |
| GPM2 | GACCCAACAGACCATAGAAACGAT | GN specific forward |
| GPM2 | CCCAACAGACCATAGAAACGAC | SB specific forward |
| GPM2 | ACAATCAGGCATGAAGATTCATCATATTGATT | qPCR reverse |
| SDH2 | CCTTACTTACAGAGATCATCGTTTCCAA | GN specific forward |
| SDH2 | CCTTACTTACAGAGATCATCGTTTCCAG | SB specific forward |
| SDH2 | ACAGACCGGGTCATAGCATTG | qPCR reverse |
| YBL036C | GTTGTAATGCTATAAAAAAAAAGATCTTCGTT | GN specific reverse |
| YBL036C | GTTGTAATGCTATAAAAAAAAAGATCTTCGTA | SB specific reverse |
| YBL036C | AGTGGCACTTTATTGGCGGT | qPCR forward |
